# MilliMap: interactive closed-loop analysis for spatial omics

**DOI:** 10.64898/2026.05.01.722104

**Authors:** Qianlu Feng, Siyuan Brant Qian, Lily Jiaxin Wan, Zachary Starr, Sarah Asif, Hee-Sun Han

## Abstract

Spatial omics analysis requires iterative interplay between statistical computation and tissue-context interpretation, yet current workflows fragment these steps across disconnected environments. We present MilliMap, an interactive framework that unifies statistical analysis with spatial exploration. By closing the analysis-visualization loop, MilliMap empowers biologists to steer parameters, refine ROIs, and validate findings within a single environment. We demonstrate its utility by delineating complex neuroanatomy and identifying niche-restricted functional states in tumor microenvironments.

## Main

Spatial transcriptomic and proteomic technologies have rapidly expanded our ability to map gene and protein expression in tissue context, enabling molecular dissection of cellular organization across development, behavior, and disease.^1–3^ These approaches generate large, high-dimensional datasets, often comprising millions of spatially resolved molecular measurements per experi-ment.^4^ Extracting biological insight from these data requires iterative interplay between spatial visualization and statistical analysis, in which hypotheses formed from tissue context are tested computationally and reinterpreted spatially.

Current workflows fragment this process across disconnected environments. Computational frame-works such as Scanpy,^5^ Squidpy,^6^ Seurat,^7^ and Giotto Suite^8^ provide powerful analytical function-ality but require programming skills and return non-interactive plots. Visualization platforms such as Vitessce,^9^ cellxgene,^10^ and TissUUmaps^11^ support interactive exploration but display precom-puted results without triggering new analysis. Consequently, users must transition between tools to define regions of interest, execute analysis, refine parameters, and interpret results (Fig. 1a). While some commercial^12, 13^ or open-source^14^ solutions provide partial integration, these approaches are typically constrained to specific platforms or limited coupling between analytical outputs and spatial representations.

**Figure 1.**
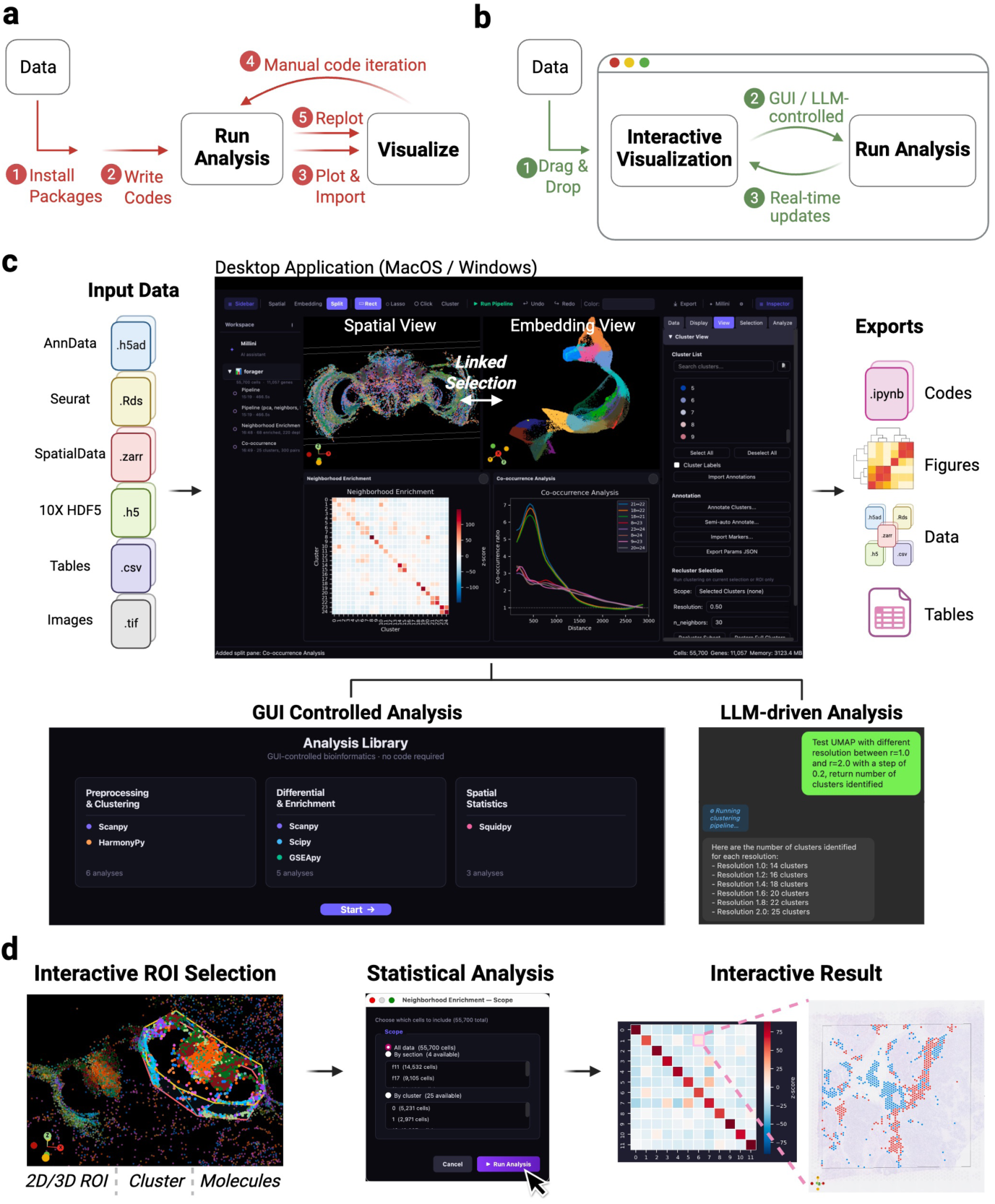
MilliMap enables interactive analysis of spatial omics data. (a) Conventional workflow. Users manually write scripts and switch to external viewers to refine parameters and test hypotheses. **(b) MilliMap workflow.** Drag-and-drop input; GUI- or LLM-driven analyses (Millini) update the linked visualization in real time. **(c)** Supported formats feed paired *Spatial* and *Embedding* views. Analysis Library and Millini outputs appear as interactive workspace cards, exportable as notebooks, figures, data, and tables. **(d)** Spatial selections (2D/3D ROIs, clusters, molecules) feed downstream analyses; interactive results update both views.

We developed MilliMap, an interactive analysis framework in which visualization and analytical computation form a continuous loop within a single environment (Fig. 1b). MilliMap links the two core capabilities required for spatial omics analysis: (i) interactive spatial and embedding views of the tissue, and (ii) a statistical analysis library built on major omics software including Scanpy,^5^ Squidpy,^6^ harmonypy,^15^ and GSEApy.^16^ These components are controlled by both graphical user interfaces (GUI) and a large language model (LLM) agent, Millini, establishing a coherent workflow for interactive analysis, with active sessions also accessible to external clients such as Claude via the Model Context Protocol (MCP) (Fig. 1c). In MilliMap’s workflow, the visual context dynamically scopes subsequent analysis, and computational outputs are returned as interactive objects that update the views in real time (Fig. 1d). MilliMap removes the technical overhead of traditional pipelines, empowering biologists to autonomously conduct in-depth analyses previously requiring dedicated computational support.

MilliMap operates across major spatial omics platforms, including Visium, Visium HD, Xenium, MERSCOPE, CosMx, and CODEX, and extends to non-spatial single-cell data (Fig. 2a–d; Extended Table 1). It takes interoperable data structures as input, including SpatialData,^17^ AnnData,^5^ and Seurat objects. MilliMap overcomes three fundamental challenges in moving from spatial measurements to biological insights.

**Figure 2.**
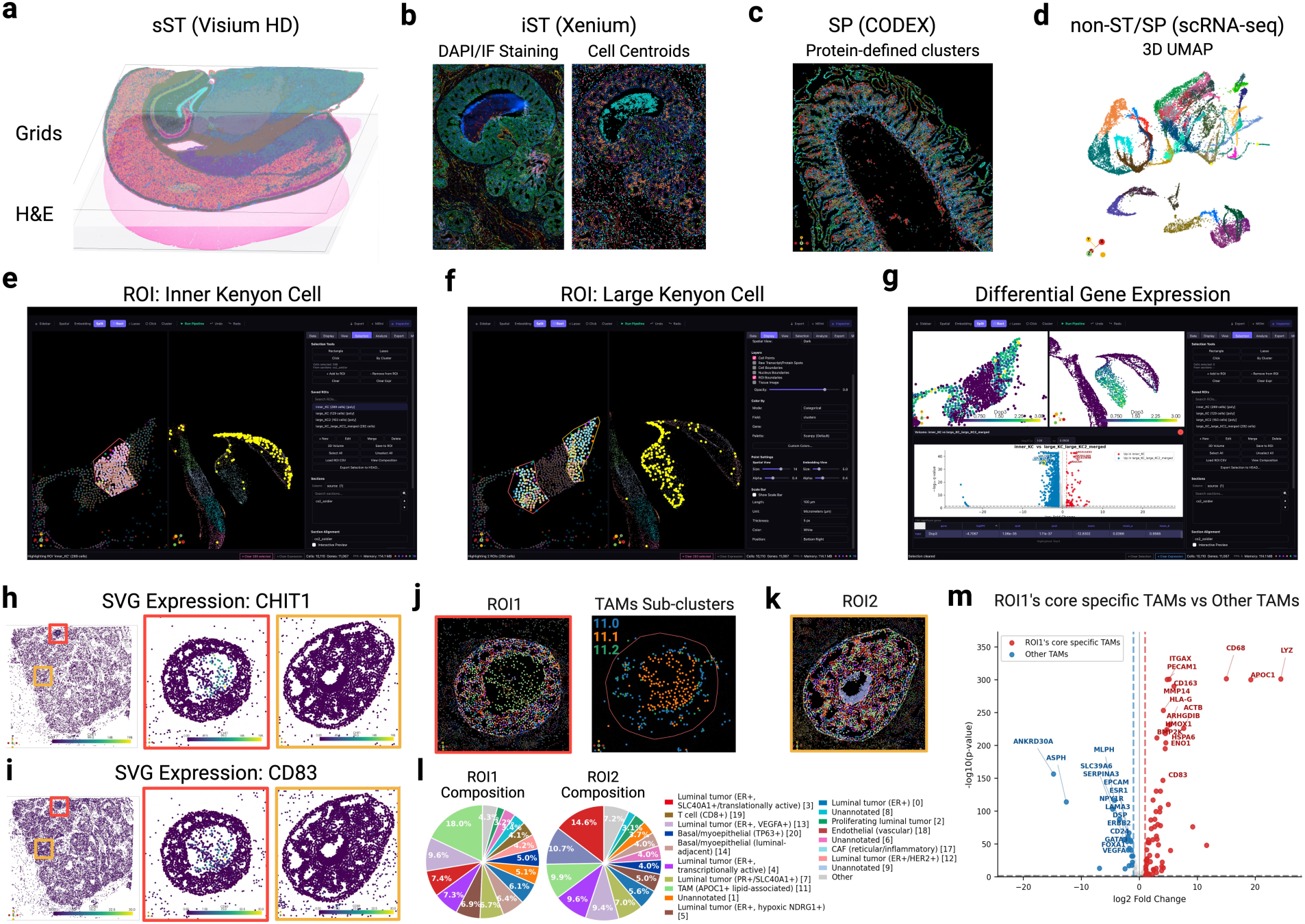
MilliMap supports diverse spatial omics data and enables biological discovery. **(a)** sST: Visium HD mouse brain, grid expression over H&E. **(b)** iST: Xenium human breast cancer; DAPI/IF morphology (left) and cluster-colored centroids (right). **(c)** SP: CODEX human intestine with protein-defined clusters. **(d)** scRNA-seq: honey bee brain, 3D UMAP. **(e, f)** Lasso-defined inner (e) and large (f) Kenyon cell (KC) ROIs (left); linked embedding confirms molecular coherence (right). **(g)** Differential expression between inner and large KCs (left: *Dop3*-colored spatial view; right: DEG heatmap). **(h)** Spatially varying gene *CHIT1* expression: whole tissue (left), ROI1 (middle), ROI2 (right). **(i)** Same layout as h, *CD83*. **(j)** Spatially resolved ROI1 cell-type clusters (left) and TAMs (cluster 11) sub-clusters (right). **(k)** Spatially resolved ROI2 cell-type clusters. **(l)** Cell type composition of ROI1 and ROI2. **(m)** Volcano of ROI1-core-specific TAMs (11.1) vs other TAMs (11.0 and 11.2).

First, MilliMap empowers high-precision, sample-specific analysis by tightly coupling parameter selection with real-time visual feedback. The spatial and embedding views are bidirectionally linked: the spatial view projects molecular features onto tissue coordinates alongside morphology images, while the embedding view (UMAP/t-SNE) captures global transcriptome- or proteome-expression structure. This synchronization allows users to tailor preprocessing, clustering, and feature selection to each tissue. For instance, users can overlay H&E or immunofluorescence images on transcript spots and iteratively refine ROIs with lasso, rectangle, or volumetric tools guided by morphology or cluster structure. Because selections in one view immediately map to the other, researchers can cross-validate selected spatial populations against expression structure before passing them to downstream analysis.

Second, MilliMap transforms the analytical process from a technical hurdle into an intuitive exten-sion of the researcher’s intent by removing the burdens of manual coding, documentation searching, and parameter interpretation. This streamlined experience is powered by a GUI-controlled analysis library that allows scientists to execute advanced workflows, including differential expression and spatial statistics on any selected subset of data, such as specific clusters or ROIs, without software configuration. Millini, an integrated LLM agent, further translates natural-language requests into multi-step workflows and provides on-the-fly explanations of parameter choices (Extended Table 2).

Third, MilliMap enhances the interpretive power of biologists by transforming static analytical outputs into live, actionable resources. Traditionally, statistical tables and figures serve as non-interactive endpoints, forcing researchers to mentally bridge numerical results by revisiting spatial architecture. MilliMap eliminates this cognitive burden by ensuring that every output functions as a dynamic driver of the interface (Fig. 1d). Selecting a transcript from a differential expression or spatially variable gene result table updates both views by expression level, providing imme-diate spatial context of candidate markers (Extended Fig. 1b–d). Selecting a cell-type pair in a neighborhood enrichment heatmap highlights both populations in the spatial view, revealing their co-localization pattern (Extended Fig. 1e). Interacting with spatial statistics plots, such as dragging a distance threshold in a co-occurrence plot, provides immediate visual feedback on proximity relationships at different scales (Extended Fig. 1f).

To ensure utility for large-scale datasets, MilliMap is engineered for both scalability and re-producibility. To maintain responsiveness for datasets exceeding millions of points, it employs adaptive level-of-detail rendering, dynamically modulating point density during navigation and restoring full resolution when the view stabilizes (Methods). On consumer-grade hardware, Mil-liMap loads datasets ranging from thousands of spots to over 4.2 million cells within seconds and sustains millisecond-level rendering latency, enabling smooth exploration without downsampling (Extended Table 3). In contrast, existing desktop and web-based visualization tools render orders of magnitude more slowly (Extended Table 4). To ensure reproducibility, MilliMap records com-plete analysis workflows and exports them as executable Jupyter notebooks, alongside result tables, figures, ROIs, and processed AnnData objects. Session archives preserve the full analytical state, allowing researchers to reopen, extend, and share findings across environments.

We demonstrate these capabilities across four datasets spanning platforms, species, and analytical tasks in Extended Figs. 1–4 (Visium, Visium HD, Xenium, CODEX), and highlight two use cases illustrating how interactive analysis enables biological discovery.

In the first use case, we delineate complex tissue boundaries where traditional domain detection lacks sensitivity. We applied a closed-loop refinement workflow to the mushroom body (MB) in a honey bee brain MERFISH dataset generated in the Han laboratory and originally reported in Lee et al.^18^ The MB is a higher-order center for learning and memory^19^ characterized by intermixed large and inner Kenyon cells (KCs) with irregular boundaries difficult to isolate by clustering alone. Existing tools make ROI definition disjointed: users inspect data in one platform (e.g., Vitessce), trace anatomical boundaries in another (e.g., Fiji), and write custom code to map regions back. Using MilliMap, ROIs are defined on tissue using morphology-guided selection and immediately projected into embedding space, where boundary cells are iteratively refined based on transcriptional structure until spatial and molecular representations are concordant (Fig. 2e,f), enabling precise separation of KC subpopulations. In-place differential expression reveals subregion-specific gene programs (Fig. 2g), including selective enrichment of *Dop3*,^20^ a D2-like dopamine receptor implicated in mushroom body learning and memory, in the large Kenyon cells.

In the second use case, we found that morphologically similar tumor nests within a single human breast tumor (10x Genomics Xenium)^21^ can harbor markedly different immune microenvironments (Fig. 2h,i). Using Moran’s *𝐼*, we identified localized *CHIT1* and *CD83* hotspots within one nest (ROI1), centered in a core surrounded by dense luminal tumor cells. Spatial profiling showed that this core was infiltrated by CD8^+^ T cells co-localized with *APOC1*^+^ lipid-associated macrophages, whereas the luminal periphery contained distinct macrophage populations (Fig. 2j,l). By con-trast, a morphologically matched control nest lacking *CHIT1*/*CD83* hotspots (ROI2) displayed an empty core and lacked both T cells and *APOC1*^+^ macrophages, while retaining similar periph-eral macrophage populations (Fig. 2k,l). To investigate these divergent niches, we performed an interactive ROI1-versus-ROI2 differential expression analysis in MilliMap. Although both nests shared luminal epithelial shells, the ROI1 periphery showed reduced type-I interferon signaling, including lower *ISG15* and *CXCL10* expression (Extended Fig. 3g), consistent with diminished chemokine cues for T-cell recruitment. These spatial patterns suggest localized immune resistance, with immune infiltration confined to the ROI1 core while the surrounding periphery may limit further recruitment. ROI2 lacked both the infiltrated core and the associated suppressive signature.

To further resolve these macrophage populations, MilliMap in-session sub-clustering identified three macrophage sub-states (11.0–11.2). The ROI1-enriched population (11.1) expressed canon-ical macrophage markers (*CD68*, *CD74*, *CD163* and *LYZ*) together with genes linked to lipid handling (*APOC1*), matrix remodeling (*MMP14*), immune inhibition (*HLA-G* and *CD83*), and anti-inflammatory maintenance (*HMOX1*) (Fig. 2j,m). Although individual components of this program have been reported in breast cancer,^22, 23^ their convergence within a spatially restricted macrophage state has not, to our knowledge, been previously described. This use case illustrates how coupled ROI selection, in-place sub-clustering and differential expression analysis can reveal highly localized cellular states^24^ that may be obscured in broader regional analyses.

MilliMap transforms spatial omics analysis from sequential pipeline execution into continuous, exploratory dialogue with the tissue. By unifying interactive visualization with high-performance statistical libraries, it keeps expert spatial intuition at the center of discovery rather than con-strained by precomputed clusters and static plots. As spatial technologies scale, MilliMap offers a platform-agnostic, interactive framework for moving from visual observation to rigorous hypothesis validation.

## Online Methods

### Software architecture and implementation

#### Desktop application design

MilliMap is a standalone Python desktop application distributed as a precompiled executable for macOS and Windows, with no dependency on external programming environments or package installations. The interface consists of two synchronized interactive viewports (a spatial coordinate view and a dimensionality-reduction embedding view), together with a tabbed control panel and a workspace panel that organizes completed analysis results (Fig. 1). Computationally intensive operations, including dimensionality reduction, clustering, and LLM-driven queries, execute in background threads to maintain interface responsiveness.

#### Data model and supported formats

All supported input formats are normalized to a unified AnnData^5^ representation upon loading.

*Spatial transcriptomics*. Supported formats include 10x Visium (Space Ranger output; H&E im-age), Visium HD (Space Ranger HD output; H&E image), 10x Xenium (Xenium Ranger output; multichannel morphology images, transcript detections, cell and nucleus boundaries), MERSCOPE (Vizgen output; H&E and staining images, transcript detections, cell boundaries), and NanoString CosMx (RNA expression, per-cell protein stain intensities, optional transcript detections). Spatial-Data (.zarr) is also supported as a generic interchange format.

*Spatial proteomics*. CODEX multiplexed imaging data (CSV; per-cell protein intensities) is sup-ported.

Generic formats including Seurat .rds and .h5ad files are accepted across both modalities. A visual column-mapping interface additionally allows users to import arbitrary tabular data with spatial coordinates by interactively assigning columns to spatial coordinates, cell identifiers, and expression values, without requiring custom parsing code. This enables compatibility with additional spatial platforms that produce a spatially annotated expression matrix.

### Analysis library

MilliMap integrates a comprehensive spatial omics analysis stack through the graphical interface with no scripting requirement. All functions are also accessible via Millini or external clients such as Claude, and are enumerated together with their underlying packages and exact versions in Extended Table 2.

#### Preprocessing and dimensionality reduction

Preprocessing follows the standard Scanpy^5^ workflow: library-size normalization and log-transformation, highly variable gene selection, optional doublet removal with Scrublet,^25^ PCA, optional Harmony^15^ batch correction, k-nearest neighbor graph construction, Leiden^26^ community detection, and UMAP^27^ or t-SNE^28^ embedding, with key parameters adjustable via GUI controls. On completion, the embedding view is populated with the computed low-dimensional layout, and both views are colored by Leiden cluster assignment.

#### Differential expression and enrichment

Differential expression between any two clusters, saved ROIs, or dataset sections uses the Wilcoxon rank-sum test (two-sided) with Benjamini–Hochberg correction (*𝑞 <* 0.05); results are stored as a paired DE table and an interactive volcano plot. Gene set enrichment queries the Enrichr API (GO Biological Process, KEGG, Reactome) via GSEApy,^16^ with additional support for pre-ranked GSEA and custom gene-set scoring; results are stored as ranked gene set tables. Cell cycle phase assignment uses the Tirosh et al. marker lists^29^ and updates both views with phase labels as a categorical colormap.

#### Spatial statistics

Spatial statistics are computed via Squidpy.^6^ Spatially variable gene detection uses Moran’s *𝐼* or Geary’s *𝐶* and produces a ranked SVG table; neighborhood enrichment returns a z-score matrix of cluster co-localization stored as an interactive heatmap; and co-occurrence analysis quantifies the probability of cluster pairs appearing within increasing distance thresholds, stored as distance-resolved curves. Per-cluster centrality scores (degree, closeness, betweenness) and cluster-pair interaction matrices further summarize the spatial neighbor graph, while Ripley’s *𝐾*, *𝐿*, and *𝐺* statistics test each cluster against complete spatial randomness. Expression-by-distance profiles compute mean gene expression as a function of distance from an anchor cell type.

### Visualization and interaction

#### Rendering and data display

MilliMap renders spatial omics data on VTK^30^ via PyVista^31^ in an off-screen OpenGL context em-bedded in the Qt interface. The spatial and embedding views share the same point-cloud pipeline, so linked selection between them is a direct array-index mapping between the two actors. Cells are drawn as interactive point clouds colored by categorical (cluster identity, cell type) or continuous (gene expression, protein intensity) colormaps. Optional overlays include segmentation polygons (Xenium, MERSCOPE) and per-gene transcript point clouds (Xenium, MERSCOPE, CosMx). Tissue morphology images, such as H&E underlays (Visium, Visium HD) and multichannel fluo-rescence stacks (Xenium, CosMx), are rendered as textured planes aligned to the spatial coordinate system. Each layer is independently toggleable with per-layer opacity sliders in the Display panel. Morphology images are lazy-loaded from OME-TIFF pyramids via tiled reads, with the active pyramid level selected to match the current zoom to keep memory use bounded. For multi-section datasets, sections are placed in a shared coordinate frame (adjustable via the Section Alignment dialog) and stacked along *𝑍*, navigated with a free 3D camera supporting orbit, pan, and zoom.

To maintain interactive frame rates on datasets exceeding a million transcripts, MilliMap imple-ments adaptive level-of-detail rendering: the transcript layer is subsampled to 500,000 points while the camera is in motion and restored to full resolution once the camera is stationary. Click-to-identify spatial picking uses vtkCellPicker against the cell point cloud.

#### Spatial selection and region of interest management

*Cell and cluster selection*. Clusters can be selected from the sidebar cluster list or defined by freeform polygon (lasso) or rectangle directly on the spatial or embedding view.

*Single-molecule selection*. Selecting a gene or protein marker from the feature list recolors all cells by its expression level as a continuous heatmap across the tissue. When raw transcript-level coordinate files are loaded, clicking a gene additionally renders each individual transcript detection as a spatially resolved point cloud, revealing the precise sub-cellular localization of that transcript.

*Region of interest (ROI) definition and use*. Any cell or transcript selection can be saved as a named ROI. For multi-section three-dimensional datasets, a volumetric selection mode allows independent polygon boundaries to be drawn on individual Z-sections; MilliMap interpolates between them to define a contiguous volumetric region spanning serial sections.

All selection types (clusters, single molecules, and saved ROIs) can be saved and directly passed as inputs to any downstream analysis, including differential expression, enrichment, and spatial statistics. When an analysis completes, both the spatial and embedding views are immediately recolored to reflect the result, enabling iterative hypothesis refinement without leaving the visual context.

#### Interactive analysis workspace

All analysis results are stored as named cards in the workspace navigator and remain accessible throughout the session. Each card opens as a result panel whose interactive behavior is tailored to the Analysis Library category that produced the result.

*Preprocessing, dimensionality reduction, and clustering*. Clustering results (Leiden) update the categorical colormap of the spatial and embedding views synchronously. Clusters can be renamed through the annotation dialog, with labels propagating instantly to all open result cards, or passed to the re-cluster function, which runs Leiden on the selected cells and returns a nested cluster track alongside the original labels. Violin plots of per-cluster gene expression support cluster-level selection: clicking a violin highlights the corresponding cells in both views.

*Differential expression*. DE tables, volcano plots, and dot plots are all gene-clickable: selecting any entry immediately recolors both views by that gene’s expression. In volcano plots, users can drag the log_2_FC and −log_10_(*𝑞*) threshold lines to restrict the highlighted gene set in real time, and clicking a volcano point additionally highlights the corresponding row in the paired DE table.

*Spatial statistics*. Spatially variable gene rankings (Moran’s *𝐼* / Geary’s *𝐶*) expose the same gene-click to spatial recolor behavior as DE tables. Clicking a cluster-pair cell in either the neighborhood enrichment heatmap or the cluster-pair interaction matrix highlights both populations on the tissue. Centrality score bar charts and Ripley’s *𝐾*, *𝐿*, and *𝐺* curves are cluster-clickable: clicking a bar or curve highlights the corresponding cluster in space. In co-occurrence plots and expression-by-distance profiles, dragging the distance threshold updates the highlighted cells in real time to show only cells within the chosen spatial range.

*Pathway enrichment and per-cell scoring*. GSEApy enrichment tables are term-clickable: selecting a term exposes its leading-edge genes, and clicking any gene recolors the spatial and embedding views by its expression. Per-cell score overlays (gene-set scoring, composite metrics) expose a color-range slider in the result card, letting users restrict the highlighted cells to those above a chosen value without re-running the analysis.

*Batch correction*. Harmony produces a corrected embedding that is selectable in the Embedding View and becomes the basis for subsequent clustering or DE analyses invoked from the Analysis Library.

Result cards can also be dragged directly onto either the spatial or embedding view as resizable floating overlay panels, enabling simultaneous display of multiple results within the same visual context.

### LLM-based analysis interfaces

#### LLM-based state-grounded analytical agent

Spatial omics analysis is inherently stateful: user requests often depend on the active dataset, its current annotations, saved ROIs, and recent analysis outputs. Millini addresses this through an LLM-based state-grounded agent powered by Google Gemini 2.5 Flash. Rather than interpreting prompts in isolation, Millini conditions each turn on the live analytical session. Here, *state* refers to a structured summary of the current session, including dataset dimensions, cluster labels, user annotations, saved ROIs, and recent differential-expression results. By injecting this dynamic context into the model’s prompt, Millini resolves session-specific entities such as “cluster 3 (Tumor)” or “ROI A” without requiring explicit re-definition, enabling iterative analytical workflows that remain strictly grounded in the evolving research state. Millini thus functions as a session-aware analytical agent rather than a generic conversational interface.

#### Tool-based orchestration of multi-step analysis

Millini does not generate free-form text or analysis scripts. Instead, it uses LLM function calling to select tools, fill parameters, and sequence analysis steps.^32, 33^ The tool registry spans the major operations supported by MilliMap, including preprocessing, clustering, differential expression, spatial statistics, enrichment analysis, and per-cell scoring (Extended Table 2). A single natural-language request can therefore trigger multiple executable analysis steps, selected iteratively as updated session state and intermediate outputs inform subsequent actions. The LLM serves only as an orchestration layer; all numerical analyses execute on the native analytical backends, shielding outputs from LLM-inherent stochasticity and hallucination.^34^ Because Millini tool calls run through the same backend routines as GUI-triggered analyses, their outputs appear in the same result cards with the same immediate spatial and embedding recoloring. Every tool call is logged with its function name, parameters, and timestamp, and feeds directly into the Jupyter notebook export path described below.^35^

#### External access via the Model Context Protocol

In addition to the in-app Millini agent, MilliMap exposes the same tool registry through an MCP server, allowing MCP-compatible clients such as Claude to invoke MilliMap analyses program-matically. The MCP server publishes the registered functions, parameter schemas, and result types used by Millini and the GUI. Tool calls execute on the identical analytical backends and surface as interactive result cards within MilliMap, with the spatial and embedding views recolored as they are for any internal call. This allows external agents to assist with parameter steering, analysis, and biological data interpretation.

### Exportable outputs and session archives

MilliMap provides three complementary export paths for reproducibility and sharing.

*Jupyter notebook export*. Every GUI- or Millini-triggered analysis is recorded in a structured session log capturing the analysis type, full parameter dictionary, and timestamp. On export, MilliMap traverses the log in chronological order and renders each entry into a parameterized Jupyter notebook cell. Per-analysis Python code templates substitute the recorded parameters and emit executable Python code with the same function signatures used internally. The resulting .ipynb contains an import cell, a data-loading cell, and one analysis cell per recorded step.

*Session archive*. A complete view of the active workspace is saved as a millimap_session_NNNN/ folder containing session.json with saved ROIs, cluster annotations, transforms, camera state, open result cards, and view settings. A companion description.json stores paths to the source dataset and morphology images. Re-opening the archive on another installation restores the full interactive state, including precomputed analytical results.

*Per-result and data exports*. Individual result cards export as tables (CSV, TSV), figures (PNG, SVG, PDF), and saved ROIs (as AnnData .obs columns). The processed dataset itself can be written to disk as .h5ad at any time, ensuring portability to any AnnData-compatible tool.

### Use cases

All use cases were analyzed using the GUI-triggered analysis pipeline described above. No external preprocessing was applied unless noted; all datasets were loaded directly into MilliMap from their respective source formats. Reproducible MilliMap session files and public dataset download instructions are provided in the usecases/ directory of the GitHub repository.

#### Extended Fig. 1: Visium human breast cancer

The Visium Human Breast Cancer dataset (Block A Section 1; 4,898 spots; 36,601 genes; Space Ranger v1.1.0) was loaded from the Space Ranger output directory. Preprocessing used default MilliMap parameters (library-size normalization, log-transformation, 2,000 highly variable genes, 50 PCs, 20 neighbors, Leiden resolution 1.0, UMAP). Differential expression used the Wilcoxon rank-sum test with Benjamini–Hochberg correction (*𝑞 <* 0.05); spatially variable genes, neigh-borhood enrichment, and co-occurrence were computed with default Squidpy parameters on the Leiden cluster key.

#### Extended Fig. 2: Visium HD mouse brain

The Visium HD Mouse Brain dataset (FFPE; C57BL/6; Space Ranger v3.0.0; 8 µm bin size; 393,543 bins; 19,059 genes) was loaded from the Space Ranger HD output directory and preprocessed with default MilliMap parameters. Canonical mouse-brain regional markers (*Gad1*, *Hpca*, *Drd2*, *Rorb*) were used for orientation. Cluster 26 was identified as hippocampal CA1/CA3 by convergence of top-ranked markers (*Hpca*, *Nrgn*, *Ppp3ca*, *Ptk2b*) and annotated via the Cluster Annotation dialog.

#### Fig. 2e–g: MERFISH honey bee brain

The MERFISH honey bee brain dataset (Han laboratory, UIUC) was loaded as an AnnData object. Two ROIs covering the inner and large Kenyon cells (KCs) of the mushroom body were constructed from morphology-guided lasso selections on the spatial view, with boundary cells iteratively refined based on transcriptional consistency in the linked embedding view. Differential expression between ROIs used the Wilcoxon rank-sum test with Benjamini–Hochberg correction (*𝑞 <* 0.05, | log_2_ FC| *>* 1).

#### Extended Fig. 3: Xenium human breast cancer

The Xenium Human Breast Biomarkers dataset^21^ (Section 1, Middle; 375,529 cells; 280-gene panel; Xenium Onboard Analysis v4.0.0) was loaded from the Xenium Ranger output directory; non-gene probes were excluded. Cells were filtered with min_counts = 20 and min_genes = 5; 308 cells in ROI2 (vs 11 in ROI1), predominantly within the acellular core, fell below threshold and were excluded from downstream analyses. Counts were CPM-normalized (1 × 10^6^) and log1p-transformed; no highly variable gene selection was applied given the 280-gene panel. PCA, k-NN graph (n_neighbors = 20), and Leiden (resolution = 1.5) yielded 21 top-level clusters, annotated against canonical breast-cancer cell-type signatures^36^ via top-ranked Wilcoxon markers; four clusters (1, 6, 8, 9) lacked markers at *𝑞 <* 0.05 and were retained as “Unannotated.” Cluster 11 (TAMs) was annotated by *CD68*/*CD163*/*CD74*/*LYZ* co-expression; M1/M2 polarization markers (*MRC1*, *ARG1*, *IL10*) were absent from the panel, so sub-states are described as functional TAM phenotypes. Cluster 11 was sub-clustered (Leiden, n_neighbors = 30, resolution = 0.5) into three sub-clusters (11.0, 11.1, 11.2); sub-cluster markers were tested for GO Biological Process enrichment via GSEApy.^16^ ROI1 and ROI2 contained 1,707 and 2,876 cells in total, respectively (inclusive of the 11 and 308 cells filtered above).

#### Extended Fig. 4: CODEX human intestine

The preprocessed and cell-type annotated CODEX Human Intestine dataset^37^ (2,603,217 cells; 53 proteins; 8 donors) was loaded. Neighborhood enrichment was computed via Squidpy with default spatial neighbor parameters on the Leiden cluster key; ranked protein markers per cluster used the Wilcoxon rank-sum test.

### Scalability and benchmarks

To scale to large datasets on consumer-grade hardware, MilliMap combines adaptive level-of-detail rendering with background-thread execution of data loading, rendering, and spatial analysis (see *Rendering and data display*). MilliMap maintains sub-5 ms mean and sub-10 ms p95 render latency up to 2.6M cells, with load times under 2 s for most datasets (Extended Table 3).

Scalability was evaluated on five real-world datasets spanning five spatial platforms and three orders of magnitude in cell count: Visium Human Breast Cancer (∼4,898 spots; 36,601 genes), CosMx SMI Human Lung (Lung5 Rep1; ∼100,292 cells; 3.7M transcripts), Xenium Human Breast Biomarkers (∼375,529 cells; 8.9M transcripts), CODEX Human Intestine (∼2,603,217 cells; 53 proteins), and MERFISH Whole Mouse Brain (WB_MERFISH_animal1_coronal; ∼4.2M cells; 1,120 genes). All values are medians of three independent runs.

For each dataset we measured four core metrics. *Load time*: wall-clock seconds from file-open to first interactive render (cell × gene matrix and spatial coordinates only; no tissue images, cell-boundary polygons, or transcripts), timed with time.perf_counter() around the platform-specific loader call. *Load memory*: increase in resident set size (RSS) across the loader call, measured via the macOS mach_task_basic_info API with resource.getrusage maxrss as fallback. *Render latency*: mean and 95th-percentile frame time during continuous pan/zoom, with a PyVista per-cluster-colored point cloud rendered off-screen at 960×800 and each frame forced through the rasterizer via camera.Modified() + ren_win.Render() (three warm-up frames discarded; 30 timed frames). *Peak memory*: maximum process RSS after all operations complete.

For datasets with per-transcript coordinate files (CosMx, Xenium), three additional metrics are reported. *Transcript load time*: seconds to read transcript coordinates from disk at 10% sampling. *Transcript render latency*: mean frame time with a transcript overlay capped at 500,000 points (matching the viewer’s LOD cap), using the same off-screen orbit protocol. *Transcript pick latency*: mean time for click-to-identify picking via 20 vtkCellPicker.Pick() calls at random jittered screen coordinates (tolerance 0.01). Platforms without individual transcript coordinates (Visium, MERFISH, CODEX) are marked with ‡.

*Cross-tool rendering benchmark (Extended Table 4)*. We compared MilliMap against napari-spatialdata,^17^ cellxgene,^10^ TissUUmaps,^11^ and Vitessce^9^ on the same datasets used above, loading an identical cluster-colored cell point cloud in each tool without transcripts, polygons, or images. We executed a fixed programmatic pan trajectory and recorded mean and 95th-percentile per-frame latency, discarding three warm-up frames and timing the next 30; reported values are medians of three runs. MilliMap and napari-spatialdata were driven through their native Python APIs; the three web tools were run in headless Chromium, which uses software GL (SwiftShader) and therefore penalizes WebGL-based renderers relative to hardware-accelerated desktop OpenGL. A dash (—) indicates the dataset is outside the tool’s supported scope.

## Acknowledgments

We thank members of the Han laboratory and Prof. Dave S. Zhao for helpful discussion. This work was supported by NIH R35GM147420 to H.-S.H.

## Author Contributions

Q.F., S.B.Q., L.J.W., and H.-S.H. conceived the idea. Q.F. designed and implemented the Mil-liMap software, including the spatial visualization engine, analysis pipeline, interactive workspace, data format support, and build and distribution infrastructure. L.J.W. designed and implemented the Millini natural-language interface and LLM function-calling backend. S.B.Q. developed performance-critical components including the cached and lazy-loading data pipeline and ren-dering optimizations. Z.S. and S.A. performed software validation, identified and reported broken logic and edge-case failures, and contributed to iterative debugging of the analysis pipeline. Q.F., H.-S.H., S.B.Q., L.J.W., and S.A. wrote the manuscript. H.-S.H. supervised the project and acquired funding.

## Data availability

All data shown in figures are available under permissive licenses. The Visium Human Breast Cancer dataset (Block A Section 1; fresh frozen invasive ductal carcinoma; Space Ranger v1.1.0) is available from 10x Genomics at https://www.10xgenomics.com/datasets/human-breast-cancer-block-a-section-1-1-standard-1-1-0, licensed under CC BY 4.0. The Visium HD Mouse Brain dataset (FFPE; C57BL/6; Space Ranger v3.0.0) is available from 10x Genomics at https://www.10xgenomics.com/datasets/visium-hd-cytassist-gene-expression-libraries-of-mouse-brain-he, licensed under CC BY 4.0. The Xenium Human Breast Biomarkers dataset^21^ (Section 1, Middle; 375,529 cells; 280-gene custom panel; Xenium Onboard Analysis v4.0.0) is available from 10x Genomics at https://www.10xgenomics.com/datasets/xenium-ffpe-human-breast-biomarkers, licensed under CC BY 4.0. The CosMx SMI FFPE Lung5 Rep1 dataset (human lung; FFPE) is available from Bruker Spatial Biology at https://brukerspatialbiology.com/resources/smi-ffpe-dataset-lung5-rep1-data/. The CODEX Human Intestine dataset^37^ (2,603,217 cells; 53 proteins; 8 donors) is available from the Dryad Digital Repository at https://doi.org/10.5061/dryad.pk0p2ngrf. The MERFISH whole mouse brain dataset used in scalability benchmarks (WB_MERFISH_animal1_coronal) is available from CZ CELLxGENE at https://cellxgene.cziscience.com/collections/0cca8620-8dee-45d0-aef5-23f032a5cf09. The MERFISH honey bee brain dataset (Fig. 2e–g) was generated in the Han laboratory at the University of Illinois Urbana-Champaign and was originally reported in Lee et al.;^18^ the raw MERFISH measurements and the scRNA-seq–imputed expression values used in this study are available from the Illinois Data Bank at https://doi.org/10.13012/B2IDB-5536668_V1. The honey bee single-cell RNA-seq dataset^19^ is available from the NCBI Gene Expression Omnibus under accession GSE142044.

## Code availability

Compiled executables for macOS and Windows are available for public use and testing at https://www.milliomics.com/millimap. The MilliMap source code and precompiled executables are currently held in a private repository. The codebase and reproducible use case sessions will be made fully public under an open-source MIT license at https://github.com/milliomics/MilliMap upon publication in a peer-reviewed journal.

---

**Extended Table 1.** Feature comparison of MilliMap against representative spatial omics analysis and visualization tools. Rows are grouped by feature category, with MilliMap’s uniquely distinguishing capabilities listed first within each group. ✓: fully supported; ×: not supported; ∼: partially supported; —: not applicable (tool is a code library, not an application).

| Feature | MilliMap | Loupe Browser<sup>12</sup> | ezSingle-Cell<sup>14</sup> | cellxgene<sup>10</sup> | Vitessce<sup>9</sup> | Squidpy<sup>6</sup> | Giotto Suite<sup>8</sup> | napari-spatialdata<sup>17</sup> | TissUU-maps<sup>11</sup> |
| --- | --- | --- | --- | --- | --- | --- | --- | --- | --- |
| *Interface and accessibility* |  |  |  |  |  |  |  |  |  |
| Code-free GUI | ✓ | ✓ | ✓ | ~ | ~ | — | — | ~ | ✓ |
| Standalone desktop app | ✓ | ✓ | × | ~ | × | — | — | ~ | ✓ |
| LLM natural-language interface | ✓ | × | × | × | × | × | × | × | × |
| *Analysis–visualization integration* |  |  |  |  |  |  |  |  |  |
| Result → spatial feedback | ✓ | ~ | × | ~ | ~ | — | — | × | ~ |
| ROI-based analysis | ✓ | ~ | × | ~ | ~ | × | ~ | × | ~ |
| 3D / multi-section | ✓ | × | × | × | ~ | × | ~ | ~ | ✓ |
| Real-time parameter steering | ✓ | ~ | × | ~ | ~ | — | — | × | ~ |
| *In-app analysis* |  |  |  |  |  |  |  |  |  |
| Preprocessing & clustering | ✓ | ~ | ✓ | ~ | × | ✓ | ✓ | ~ | × |
| Differential expression | ✓ | ✓ | ✓ | ~ | × | ✓ | ✓ | ~ | × |
| Spatial statistics | ✓ | × | ~ | × | × | ✓ | ✓ | ~ | × |
| Pathway enrichment | ✓ | × | ~ | × | × | × | ~ | × | × |
| *Reproducibility and platform support* |  |  |  |  |  |  |  |  |  |
| Jupyter notebook export | ✓ | × | × | × | × | — | — | — | × |
| Session export (analysis + visualization state) | ✓ | ~ | × | × | ~ | — | — | — | ~ |

**Extended Table 2.**
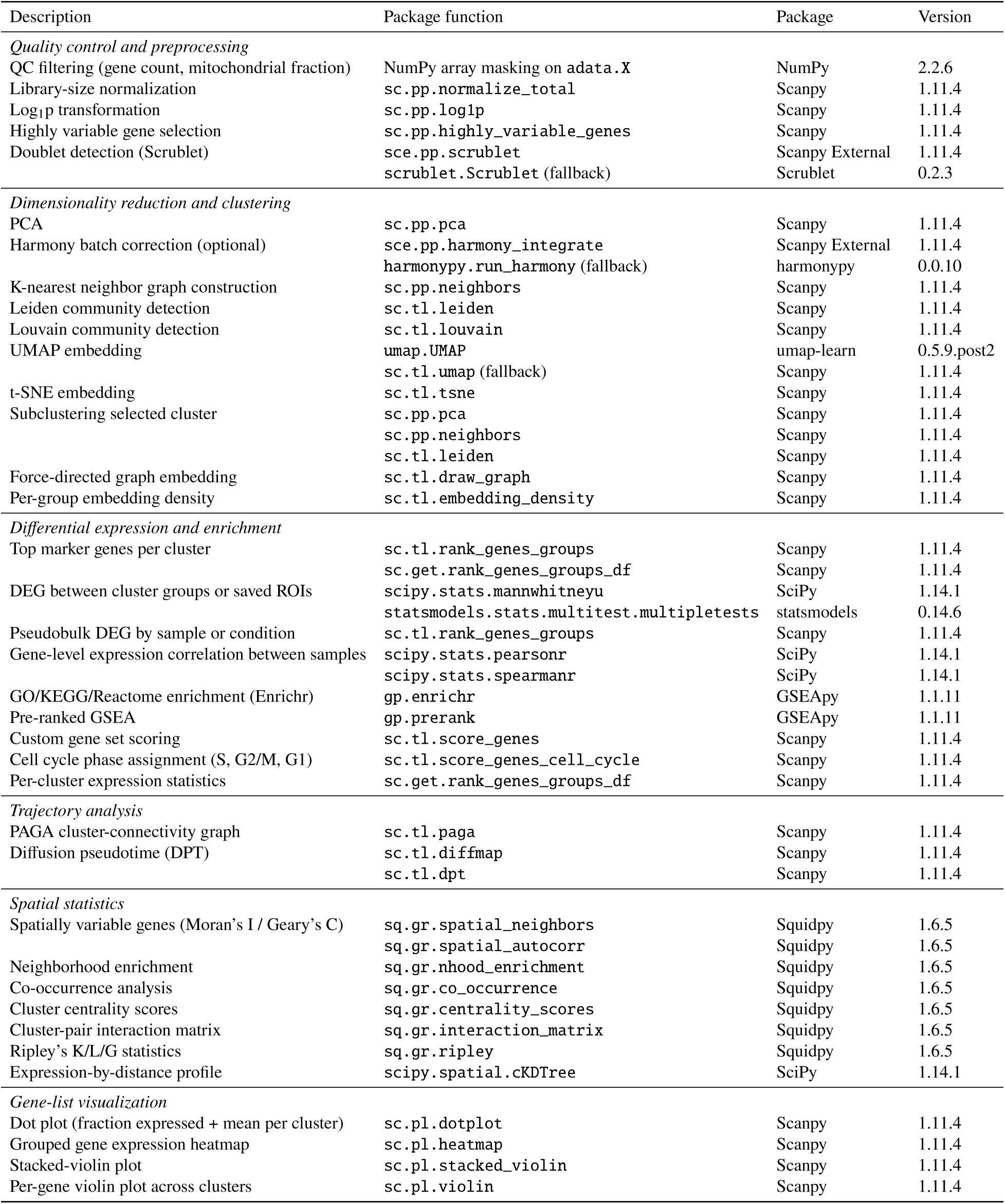
Package functions called by MilliMap analyses, with the package version used. Each row corresponds to a single function call; analyses that invoke multiple functions appear on consecutive rows.

| Description | Package function | Package | Version |
| --- | --- | --- | --- |
| *Quality control and preprocessing* |  |  |  |
| QC filtering (gene count, mitochondrial fraction) | NumPy array masking on adata.X | NumPy | 2.2.6 |
| Library-size normalization | sc.pp.normalize_total | Scanpy | 1.11.4 |
| Log<sub>1</sub>p transformation | sc.pp.log1p | Scanpy | 1.11.4 |
| Highly variable gene selection | sc.pp.highly_variable_genes | Scanpy | 1.11.4 |
| Doublet detection (Scrublet) | sce.pp.scrublet | Scanpy External | 1.11.4 |
|  | scrublet.Scrublet (fallback) | Scrublet | 0.2.3 |
| *Dimensionality reduction and clustering* |  |  |  |
| PCA | sc.pp.pca | Scanpy | 1.11.4 |
| Harmony batch correction (optional) | sce.pp.harmony_integrate | Scanpy External | 1.11.4 |
|  | harmonypy.run_harmony (fallback) | harmonypy | 0.0.10 |
| K-nearest neighbor graph construction | sc.pp.neighbors | Scanpy | 1.11.4 |
| Leiden community detection | sc.tl.leiden | Scanpy | 1.11.4 |
| Louvain community detection | sc.tl.louvain | Scanpy | 1.11.4 |
| UMAP embedding | umap.UMAP | umap-learn | 0.5.9.post2 |
|  | sc.tl.umap (fallback) | Scanpy | 1.11.4 |
| t-SNE embedding | sc.tl.tsne | Scanpy | 1.11.4 |
| Subclustering selected cluster | sc.pp.pca | Scanpy | 1.11.4 |
|  | sc.pp.neighbors | Scanpy | 1.11.4 |
|  | sc.tl.leiden | Scanpy | 1.11.4 |
| Force-directed graph embedding | sc.tl.draw_graph | Scanpy | 1.11.4 |
| Per-group embedding density | sc.tl.embedding_density | Scanpy | 1.11.4 |
| *Differential expression and enrichment* |  |  |  |
| Top marker genes per cluster | sc.tl.rank_genes_groups | Scanpy | 1.11.4 |
|  | sc.get.rank_genes_groups_df | Scanpy | 1.11.4 |
| DEG between cluster groups or saved ROIs | scipy.stats.mannwhitneyu | SciPy | 1.14.1 |
|  | statsmodels.stats.multitest.multipletests | statsmodels | 0.14.6 |
| Pseudobulk DEG by sample or condition | sc.tl.rank_genes_groups | Scanpy | 1.11.4 |
| Gene-level expression correlation between samples | scipy.stats.pearsonr | SciPy | 1.14.1 |
|  | scipy.stats.spearmanr | SciPy | 1.14.1 |
| GO/KEGG/Reactome enrichment (Enrichr) | gp.enrichr | GSEAPy | 1.1.11 |
| Pre-ranked GSEA | gp.prerank | GSEAPy | 1.1.11 |
| Custom gene set scoring | sc.tl.score_genes | Scanpy | 1.11.4 |
| Cell cycle phase assignment (S, G2/M, G1) | sc.tl.score_genes_cell_cycle | Scanpy | 1.11.4 |
| Per-cluster expression statistics | sc.get.rank_genes_groups_df | Scanpy | 1.11.4 |
| *Trajectory analysis* |  |  |  |
| PAGA cluster-connectivity graph | sc.tl.paga | Scanpy | 1.11.4 |
| Diffusion pseudotime (DPT) | sc.tl.diffmap | Scanpy | 1.11.4 |
|  | sc.tl.dpt | Scanpy | 1.11.4 |
| *Spatial statistics* |  |  |  |
| Spatially variable genes (Moran's I / Geary's C) | sq.gr.spatial_neighbors | Squidpy | 1.6.5 |
|  | sq.gr.spatial_autocorr | Squidpy | 1.6.5 |
| Neighborhood enrichment | sq.gr.nhood_enrichment | Squidpy | 1.6.5 |
| Co-occurrence analysis | sq.gr.co_occurrence | Squidpy | 1.6.5 |
| Cluster centrality scores | sq.gr.centrality_scores | Squidpy | 1.6.5 |
| Cluster-pair interaction matrix | sq.gr.interaction_matrix | Squidpy | 1.6.5 |
| Ripley's K/L/G statistics | sq.gr.ripley | Squidpy | 1.6.5 |
| Expression-by-distance profile | scipy.spatial.cKDTree | SciPy | 1.14.1 |
| *Gene-list visualization* |  |  |  |
| Dot plot (fraction expressed + mean per cluster) | sc.pl.dotplot | Scanpy | 1.11.4 |
| Grouped gene expression heatmap | sc.pl.heatmap | Scanpy | 1.11.4 |
| Stacked-violin plot | sc.pl.stacked_violin | Scanpy | 1.11.4 |
| Per-gene violin plot across clusters | sc.pl.violin | Scanpy | 1.11.4 |

**Extended Table 3.** MilliMap scalability benchmarks across five spatial omics datasets spanning five platforms (Visium, CosMx, MERFISH, Xenium, CODEX) and three orders of magnitude in cell count (4.9K–4.2M). *Load time*: seconds from file-open to first interactive render (cell × gene matrix and spatial coordinates only; no tissue images or cell-boundary polygons). *Render mean / p95*: mean and 95th-percentile frame render latency (ms) during continuous pan/zoom. *Peak memory*: maximum OS process resident set size (RSS). *Transcript load time*: seconds to read per-transcript coordinates from disk (separate from dataset open). *Transcript render mean*: mean frame render latency (ms) with the transcript point-cloud overlay active. *Transcript pick mean*: mean latency (ms) for spatial picking (click-to-identify) on the transcript layer. ‡Dataset does not provide individual transcript coordinates.

| Metric | Visium<br>Breast Cancer<br>(~4.9K spots<br>36,601 genes) | CosMx<br>Lung Rep1<br>(~100K cells<br>3.7M transcripts) | MERFISH<br>Whole Mouse Brain<br>(~4.2M cells<br>1,120 genes) | Xenium<br>Human Breast<br>(~376K cells<br>8.9M transcripts) | CODEX<br>Intestine<br>(~2.6M cells<br>53 proteins) |
| --- | --- | --- | --- | --- | --- |
| *Visualization (load time, interactivity, memory)* |  |  |  |  |  |
| Load time (s) | 1.73 | 1.73 | 25.30 | 0.36 | 6.03 |
| Render mean (ms) | 0.84 | 0.81 | 17.94 | 1.17 | 4.64 |
| Render p95 (ms) | 2.78 | 2.39 | 25.77 | 2.63 | 9.11 |
| Peak memory (GB) | 1.2 | 4.0 | 11.6 | 4.3 | 6.6 |
| *Transcript-level rendering (Xenium & CosMx only)* |  |  |  |  |  |
| Transcript load<br>time (s)‡ | ‡ | 1.14 | ‡ | 0.59 | ‡ |
| Transcript render<br>mean (ms)‡ | ‡ | 0.95 | ‡ | 1.12 | ‡ |
| Transcript pick<br>mean (ms)‡ | ‡ | 9.0 | ‡ | 8.85 | ‡ |

**Extended Table 4.** Interactive rendering benchmarks for MilliMap against representative spatial omics visualization tools across four real-world datasets spanning three orders of magnitude in cell count. Each cell reports mean and 95th-percentile (p95) frame latency during programmatic pan of a cluster-colored point cloud; lower is better. Mean reflects typical frame time; p95 reflects realistic worst-case latency (95% of frames render at or below this value). ^a^ Web tools (cellxgene, TissUUmaps, Vitessce) were rendered in headless Chromium with software GL (SwiftShader), which penalizes WebGL-based renderers relative to hardware-accelerated desktop GL. —: dataset not within the supported scope of that tool (e.g., unsupported data type or scale). All values are medians over three independent runs with the first three warm-up frames discarded.

| Tool | Visium | Xenium | CODEX | MERFISH |
| --- | --- | --- | --- | --- |
|  | Human Breast<br>(~4.9K spots) | Human Breast<br>(~376K cells) | Human Intestine<br>(~2.6M cells) | Whole Mouse Brain<br>(~4.2M cells) |
| *Mean / p95 frame latency (ms)* |  |  |  |  |
| MilliMap | **0.8 / 2.8** | **1.2 / 2.6** | **4.6 / 9.1** | **17.9 / 25.8** |
| napari-spatialdata | 2.8 / 3.1 | 6.2 / 6.8 | — | — |
| cellxgene<sup>a</sup> | 8.5 / 9.9 | 82.4 / 97.1 | — | — |
| TissUUmaps<sup>a</sup> | 8.4 / 9.5 | 120.0 / 127.4 | 799.3 / 833.0 | 1033.9 / 1160.0 |
| Vitessce<sup>a</sup> | 60.3 / 68.8 | 4323.9 / 5063.5 | — | — |

**Extended Fig. 1.**
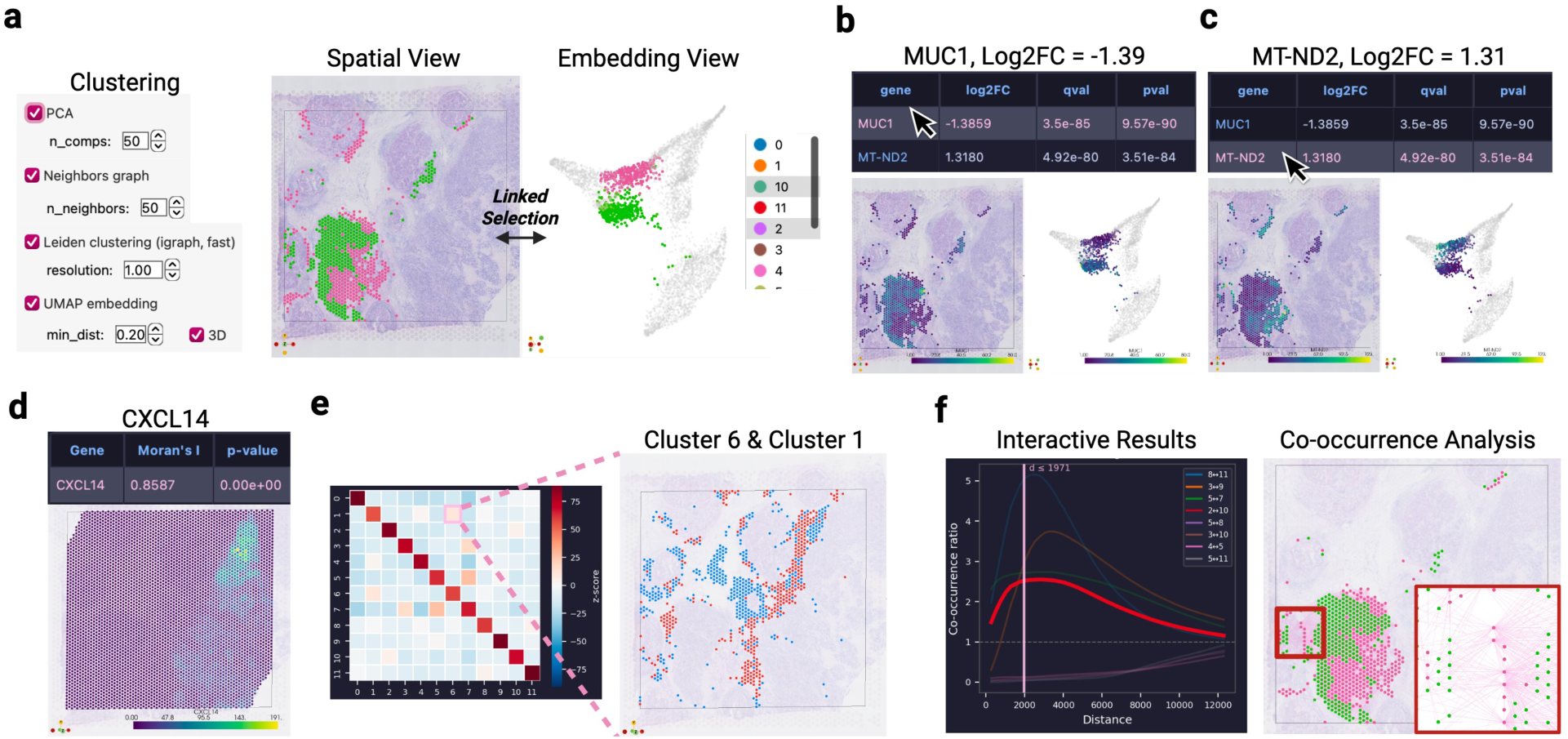
Visium human breast cancer: code-free analysis and interactive results. **(a)** Code-free preprocessing and clustering (PCA, neighbor graph, Leiden, UMAP). Spatial view (left) and linked UMAP (right) colored by Leiden cluster; lasso selection in one view highlights the same cell clusters in the other. **(b)** Interactive differential expression: selecting *MUC1* in the DEG table recolors both views by expression. **(c)** Interactive differential expression: selecting *MT-ND2* in the DEG table recolors both views by expression. **(d)** Interactive spatially variable gene discovery: selecting *CXCL14* in the Moran’s *𝐼* table recolors the spatial view. **(e)** Interactive neighborhood enrichment: selecting a cluster pair in the z-score heatmap highlights both populations in space. **(f)** Interactive co-occurrence: selecting a cluster-pair curve highlights the populations in space; dragging the distance threshold updates the highlighted cells in real time.

**Extended Fig. 2.**
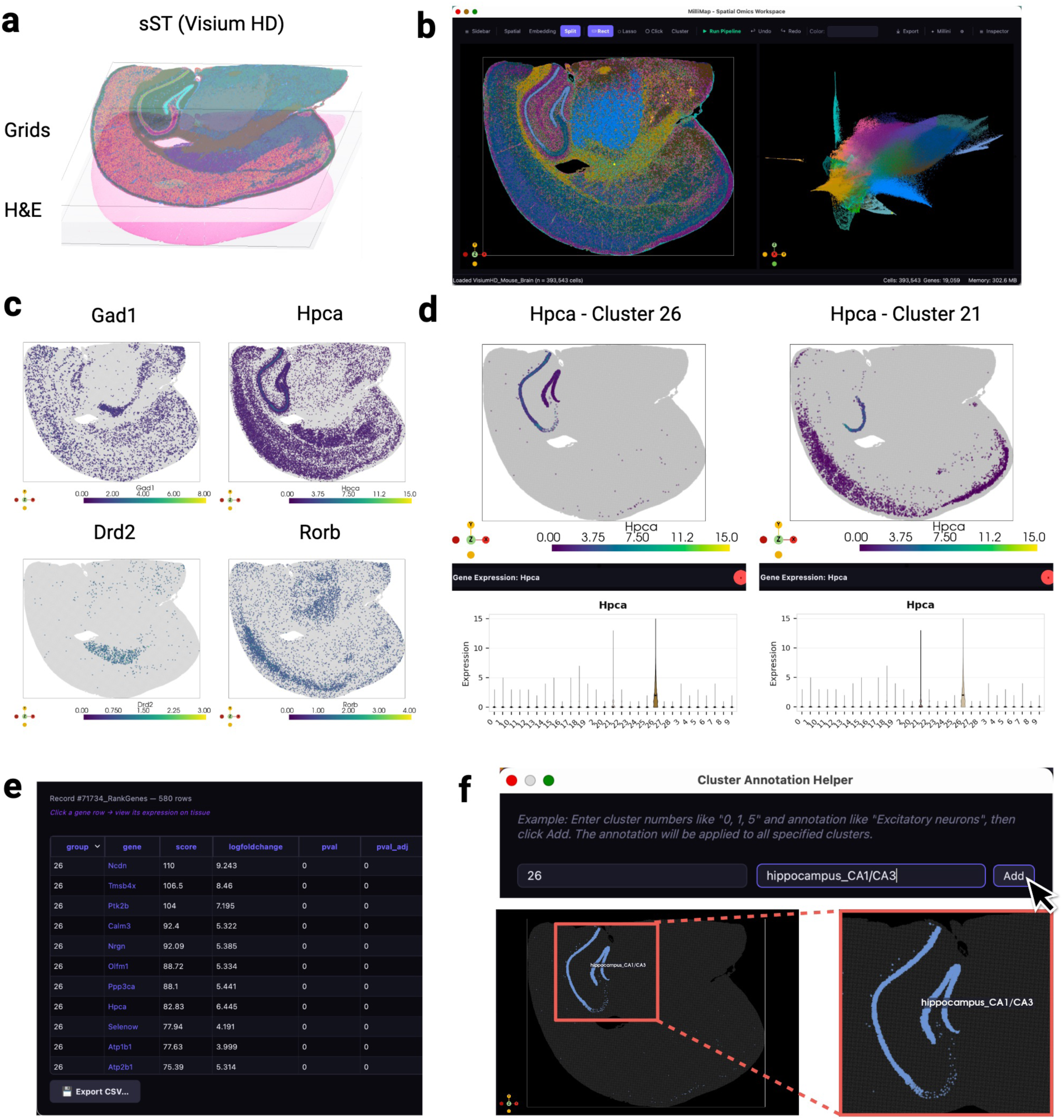
Visium HD mouse brain: clustering, marker exploration, and region annotation. **(a)** 3D spatial view of the Visium HD dataset (8 µm binning) with H&E and grid overlay. **(b)** Leiden clustering (left) and linked UMAP (right) after preprocessing (393,543 bins; 19,059 genes). **(c)** Spatial expression of *Gad1*, *Hpca*, *Drd2*, and *Rorb*. **(d)** Cluster-specific *Hpca* expression: cluster 26 (left) vs. cluster 21 (right). **(e)** Top marker genes for cluster 26. **(f)** Cluster 26 is annotated as hippocampus_CA1/CA3 via the annotation dialog.

**Extended Fig. 3.**
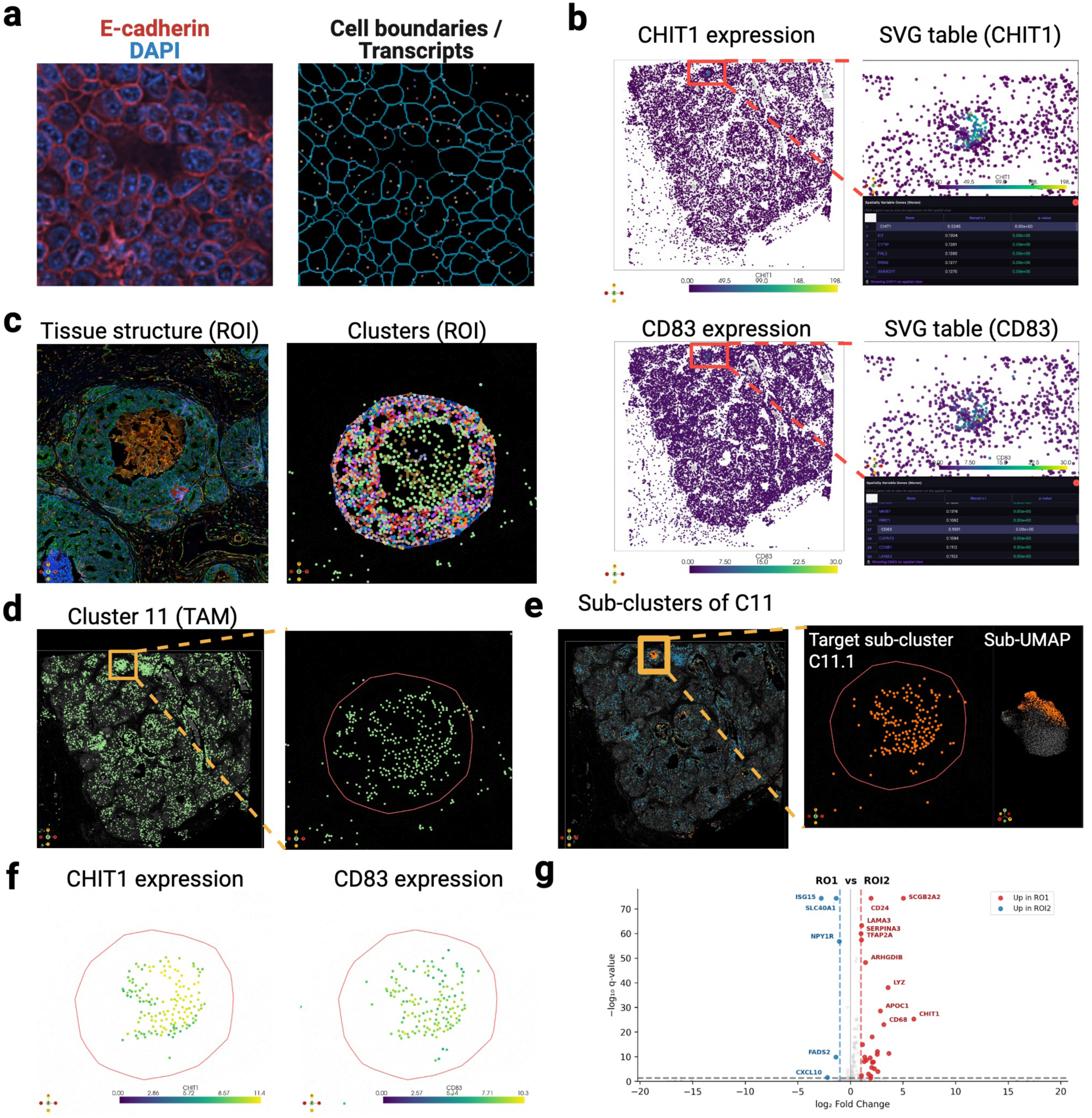
Xenium subcellular analysis: SVG-guided sub-clustering and functional annotation. **(a)** Xenium data in MilliMap: multichannel fluorescence (E-cadherin, red; DAPI, blue; left) and cell polygons with transcript detections (right). **(b)** Spatially variable gene discovery (Moran’s *𝐼*): *CHIT1* and *CD83* show regional enrichment; interactive gene selection in the SVG ranking table (right) recolors the spatial view in place. **(c)** ROI inspection: Leiden cluster overlay (left) and zoomed cluster architecture in the ROI1 core (right). **(d)** Cluster 11 (TAMs), the most enriched cluster near the SVG-high compartment, is spatially diffuse. **(e)** Sub-clustering of Cluster 11 (TAMs) reveals a confined subpopulation (sub-cluster 11.1) in the *CHIT1*/*CD83*-enriched region (left: spatial view; right: sub-UMAP). **(f)** *CHIT1* and *CD83* expression in the Cluster 11 (TAM) sub-cluster (spatial view). **(g)** Volcano plot of differential expression between ROI1 and ROI2, showing molecular divergence between the two morphologically matched tumor nests.

**Extended Fig. 4.**
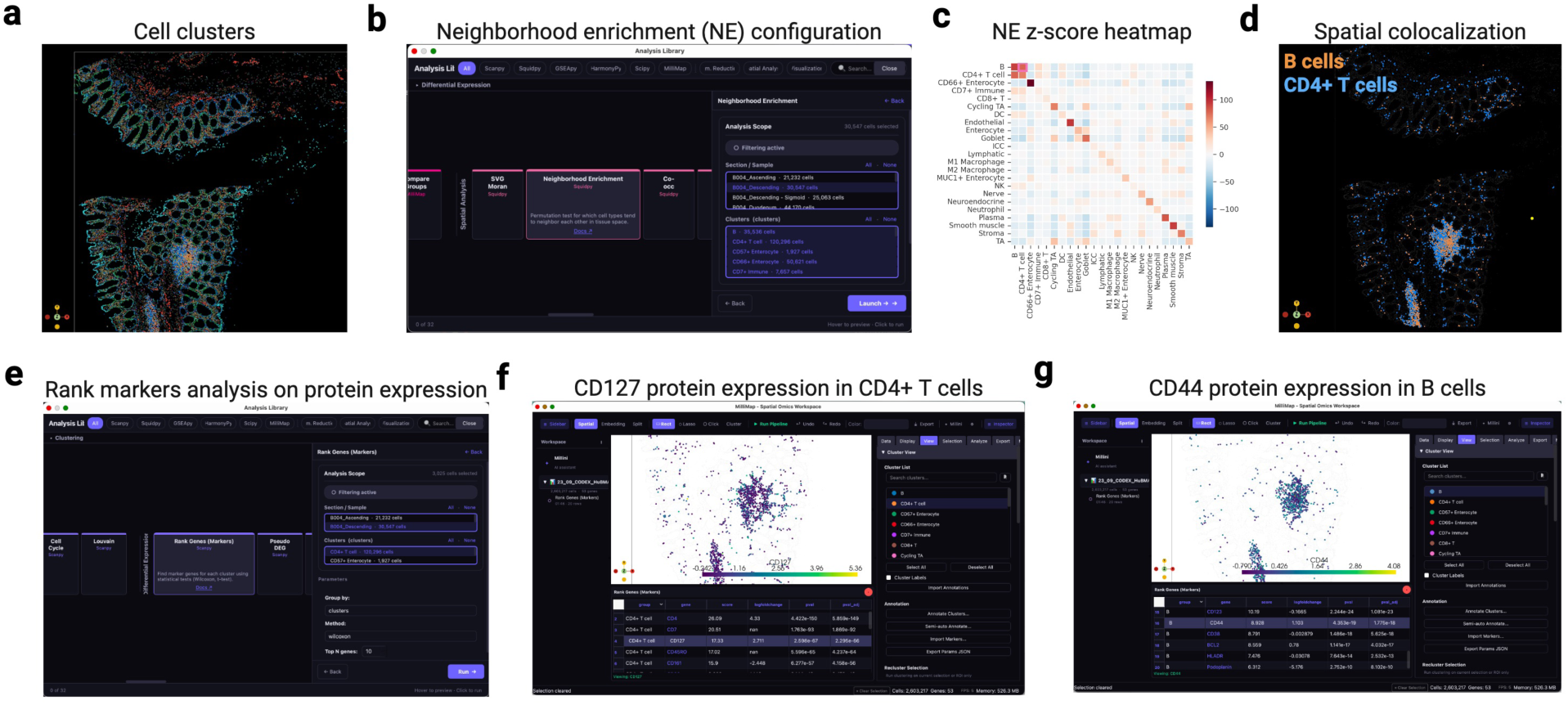
CODEX human intestine: neighborhood enrichment and protein marker exploration. **(a)** Spatial view of the CODEX dataset (2.6M cells; 53 proteins) colored by Leiden clusters. **(b)** Neighborhood enrichment parameter panel. **(c)** Enrichment z-score heatmap: red, co-localization; blue, avoidance. **(d)** Clicking the B cell / CD4^+^ T cell pair highlights both populations in space. **(e)** Ranked protein markers per cluster with score, log fold change, and adjusted *𝑞*-value. **(f)** CD127 expression in CD4^+^ T cells. **(g)** CD44 expression in B cells.

